# Towards improving full-length ribosome density prediction by bridging sequence and graph-based representations

**DOI:** 10.1101/2024.04.08.588507

**Authors:** Mohan Vamsi Nallapareddy, Francesco Craighero, Cédric Gobet, Felix Naef, Pierre Vandergheynst

## Abstract

Translation elongation plays an important role in regulating protein concentrations in the cell, and dysregulation of this process has been linked to several human diseases. In this study, we use data from ribo-seq experiments to model ribosome densities, and in turn, predict the speed of translation. The proposed method, RiboGL, combines graph and recurrent neural networks to account for both graph and sequence-based features. The model takes a graph representing the secondary structure of the mRNA sequence as input, which incorporates both sequence and structural codon neighbors. In our experiments, RiboGL greatly outperforms the state-of-the-art RiboMIMO model for ribosome density prediction. We also conduct ablation studies to justify the design choices made in building the pipeline. Additionally, we use gradient-based interpretability to understand how the codon context and the structural neighbors affect the ribosome density at the A-site. By individually analyzing the genes in the dataset, we elucidate how structural neighbors could also potentially play a role in defining the ribosome density. Importantly, since these neighbors can be far away in the sequence, a recurrent model alone could not easily extract this information. This study lays the foundation for understanding how the mRNA secondary structure can be exploited for ribosome density prediction, and how in the future other graph modalities such as features from the nascent polypeptide can be used to further our understanding of translation in general.

## 1 Introduction

Translation, the process in which RNA nucleotide triplets are encoded into amino acids to build proteins, plays a vital role in cell function. Translational control allows for rapid changes in the concentrations of encoded proteins in the cells. Thus, translational control plays an important role in maintaining homeostasis, and in modulating more permanent changes in cell fate or physiology [22]. Errors in the translation machinery are the cause of a variety of human diseases, including certain cancers and metabolic disorders. Dysregulation of signaling pathways that control cell growth and proliferation can lead to cancers, and these pathways also affect translation. Cancer is also associated with abnormal changes in the amounts of tRNAs, translation regulatory factors, and initiation factors. In particular, translation elongation has emerged as an important process that is often dysregulated in these diseases [8].

Ribo-seq is a technique to obtain ribosome read count values at codon resolution. Learning how one can predict these values can help us understand important translation-related phenomena such as ribosome stalling, ribosome collisions, and synonymous codon bias. This could also potentially elucidate the underlying mechanisms of various metabolic diseases. Recently, we have seen the application of machine learning models to predict ribosome read count values using information from the mRNA sequence [14, 17, 29, 25, 11, 24].

In this study, we model the mRNA sequence as a graph using its predicted secondary structure and design a graph-based approach to predict full-length ribosome densities (see **Figure 1**). To the best of our knowledge, the proposed approach, named RiboGL, is the first one that leverages the graph nature of the mRNA secondary structure. This approach is different from the ones proposed in the literature, where the mRNA is modeled in terms of a sequence [24] or as a codon context window [29]. More in detail, the mRNA is encoded as a graph with edges between neighboring codons, along with edges between structural neighbors which are derived from the secondary structure. Additionally, RiboGL is the first one to learn the embeddings of the codons, as opposed to using one-hot encodings as the features.

**Figure 1.**
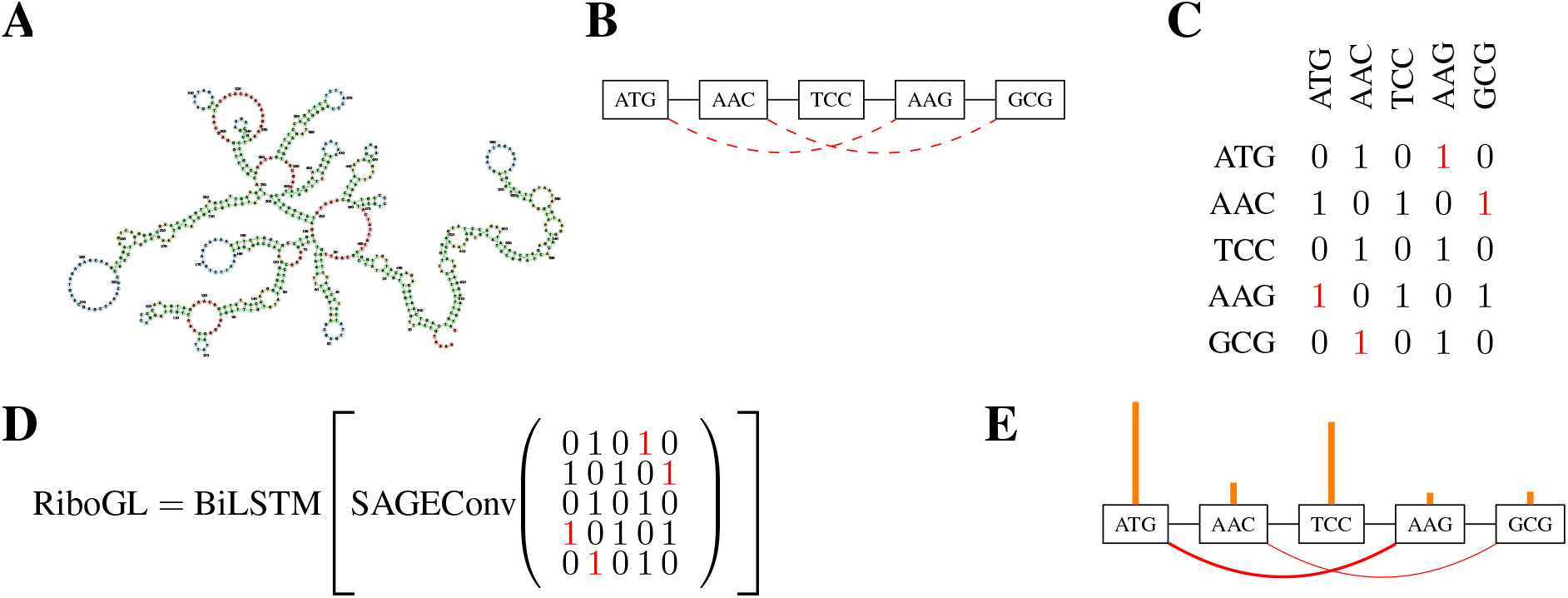
Overview of the RiboGL pipeline. (A) The mRNA secondary structure of the example gene “Wdr43” as predicted by the ViennaRNA web server [15]. (B) Graphical depiction of how the edges are modeled using an example sequence. The two kinds of edges, sequence and structural, are depicted using two colors, black and red respectively. (C) The adjacency matrix of the example graph is outlined in (B), with structural edges in red. (D) The graph processing equation for the example sequence. Firstly, a SAGE Convolution operation is applied on the input graph, and then a BiLSTM is applied to the learned node features. (E) Interpretability analysis on the example sequence - this can help understand the attributions of all the codons and the edges in the graph. The height of the orange bars indicates the magnitude of contribution from the codons, and the intensity of red on the edges indicates the magnitude of contribution from the edges.

Previous approaches for predicting the ribosome density at a specific position exploited only the information of neighboring codons [24]. By taking into account the secondary structure, the model can exploit structural neighbors to capture codons at a much greater distance. Indeed, learning such interactions with a small and noisy dataset might be prohibitive. This is also particularly relevant since sequence-based models like Recurrent Neural Networks (RNNs) might fail in capturing long-term dependencies due to vanishing gradients [18]. Conversely, Graph Neural Networks (GNNs) can easily handle the structural neighbors by accepting the secondary structure as input, but might fail at capturing information from distant nodes [1]. Consequently, we defined RiboGL as a GNN graph processing block, to extract the information from the structural neighbors, followed by a sequence processing block, overcoming the limitations of GNNs. For the graph convolution algorithm, we employed SAGE Convolutions (SAGEConv) [9], which in our experiments reached one of the highest performances with a relatively lower number of parameters. The node features learned by the GNN are then processed by the sequence block composed of a Bi-Directional Long Short Term Memory (LSTM) [10] model. **Figure 1** outlines the summarized RiboGL pipeline.

Modeling ribosome density prediction with both sequence and graph features allows us to learn new codon interactions, that can be extracted with post-hoc interpretability techniques [20]. Indeed, while with sequence-based models we can estimate only codon-wise feature importance, graph inputs enable us to extract the flow of information between neighbors through edge importance.

### Contributions

In section 4, we show that the proposed RiboGL model outperforms the state-of-the-art RiboMIMO [24] model by ∼ 21%. In section 4.1 we conduct ablation studies to justify our design choices for RiboGL. Lastly, in sections 4.2 and 4.3, we show the applications of RiboGL for interpretability, where we extract the contribution of both the codons and the edges of the secondary structure graph. To reproduce the experiments and explore predictions and attributions of RiboGL on a Mouse Liver Dataset (introduced in 3.1) refer to our GitHub repository ^2^.

## 2 Related Work on Ribosome Density Prediction

Earlier approaches to predict ribosome density, namely riboShape [14] and RUST [17], employed linear models and codon-wise statistics to infer ribo-seq densities. To denoise the dwell times, the former employed wavelets and kernel smoothing, while the latter binarized values to reduce the impact of outliers. More recent approaches exploited deep learning to handle the complexity of the ribo-seq data. ROSE [29] trained a Convolutional Neural Network (CNN) as a binary classifier to detect stalling events. Iχnos [25] and Riboexp [11] used a Feedforward and a Recurrent Neural Network (FNN and RNN), respectively, to predict each codon density using both the information about its neighbors and their RNA folding energy. While previous approaches took into consideration a window or context around the target codon, the current state-of-the-art, RiboMIMO [24], is trained to predict the whole density profile of a transcript. Similar to ROSE, RiboMIMO considers a simplified classification task to predict dwell times, while also adding another regression loss on the normalized counts. In **Table 1**, we summarized the characteristics of each approach.

**Table 1:** Summary of ribosome density modeling approaches. <u>Features</u>: codon/nucleotide/aminoacid of the A-site context (ctx) or the full sequence (seq), mRNA folding energies (fold) and secondary structure (graph). <u>Label Correction</u>: denoising (denoise), quantization (quant), and normalization, e.g., dividing by average transcript density (norm). <u>Output</u>: at codon (cdn) or sequence (seq) level. Predictor: Codon-wise Statistics (CS), Linear Model (LM), and Feedforward, Convolutional and Recurrent Neural Network (FNN, CNN, and RNN, respectively).

| Method | Features | Label Corr. | Out | Predictor |
| --- | --- | --- | --- | --- |
| riboShape [14] | ctx | denoise | cdn | LM |
| RUST [17] | ctx | quant | cdn | CS |
| ROSE [29] | ctx | quant | cdn | CNN |
| Ixnos [25] | ctx, fold | norm | cdn | FNN |
| Riboexp [11] | ctx, fold | norm | cdn | RNN |
| RiboMIMO [24] | seq, fold | norm, quant | seq | RNN |
| RiboGL (ours) | seq, graph | norm | seq | GNN+RNN |

## 3 Methods

### 3.1 Mouse Liver Dataset

Mouse liver ribosome profiling data from [4], available in the Gene Expression Omnibus (GEO) database under accession number GSE73553, was used in this study. This data was pre-processed through the initial steps of our “Ribo-DT” snakemake pipeline^3^, with slight modifications, to generate position-specific ribosome A-site coordinates on the transcriptome, as outlined in [7]. Specifically, mouse genome sequences (GRCm38/mm10), and transcript annotations were downloaded from ENSEMBL (Release 95). Sequence Read Archive (SRA) files were retrieved using the GEO accession number and then converted into FASTQ format. These files were then aligned to the mouse genome using STAR with inline adapter clipping (TGGAATTCTCGGGTGCCAAGG). The resulting BAM files were indexed. Size-dependent A-site positions were computed using a pile-up of 5’-end read density at the start codon for each read size and frame. Unique mapping reads of size between 26 and 35 nucleotides and up to one mismatch were included in the analysis. Read counts and coding DNA sequence (CDS) positions were retrieved, with the A-site offset adjusted accordingly. The codon-level ribosome counts were normalized by the gene average, similar to previous approaches [24]. Moreover, we only kept sequences with coverage, defined as the percentage of non-zero and non-NaN annotations, greater than 30%. The ranges in the annotations were reduced by applying a log1p function. The resulting dataset consisted of 6,988 genes, with 20% left out as a test set. More details on our preprocessing are available in Appendix A.1, while the relationship between coverage density and sequence length have been reported in **Figures A.1 and A.2**.

### 3.2 Secondary Structure Prediction

The secondary structure of the mRNA sequence was used as the input to the RiboGL model. The graphs of the mRNA sequences were obtained by using the ViennaRNA [15] module with the Minimum Free Energy (MFE) approach [30] (`RNA.fold` from the ViennaRNA Python API). An example structure is reported in **Figure 1 (A)**. This module results in the dot-bracket secondary structure for the mRNA, which is then converted into an adjacency matrix. The created adjacency matrix would correspond to the nucleotides, this is then pooled to create a codon-level adjacency matrix which is used as the input to the RiboGL model. An example sequence and the corresponding adjacency matrix have been mentioned in **Figures 1 (B) and (C)**.

### 3.3 Graph Neural Networks (GNNs)

Graph Neural Networks are a special class of neural networks designed to process graph-based inputs. Consider a graph *G* = (*V, E*) where node set *V* represents the *n* nodes and edge set *E* represents the *m* edges. The adjacency matrix *A* ∈ (0, 1)^*n×n*^ is populated based on the edges. For an undirected edge between nodes *i* and *j*, the values of *a*_*i,j*_ and *a*_*j,i*_ are both set to 1. GNNs follow a Message Passing Neural Network (MPNN) paradigm where the information from neighboring nodes is used to compute new node-level features in every pass of the network. The node features of node *i* are represented as *x*_*i*_. The *k*^*th*^ pass of the MPNN, is defined by aggregation functions (*AGG*^(*k*)^) and combination functions (*COM* ^(*k*)^) which are used iteratively to compute embeddings 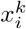 for node *i* based on messages 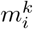 that contain information from its neighbours.

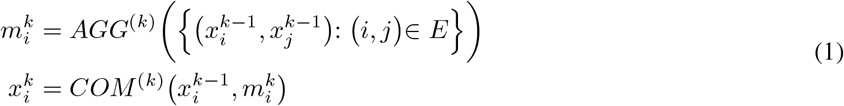

The general working of the MPNN has been outlined using equation 1. The *AGG*^(*k*)^ and *COM* ^(*k*)^ vary with different implementations of the MPNN and can result in different architectures such as the Graph Convolutional Network (GCN) [12], and the Graph Attention Network (GAT) [26].

In this study, we intend to explore how the neighborhood of the codon A-site affects the ribosome density. Studying the influence of all the edges and nodes in the graph would help us understand more about the features of the codon neighborhood that affect the speed of translation. There have been several graph interpretability algorithms that have been suggested previously [28], but to the best of our knowledge, they have not been applied to the setting of ribosome density prediction.

### 3.4 RiboGL

The RiboGL model takes as input the embedding of the gene sequence at the codon level along with random walk positional encodings [6] generated from the secondary structure graph. The model consists of two blocks in series:

1. **Graph Processing Block (GNN):** The purpose of the graph processing block is to exploit the graph structure of the mRNA, and it consists of four graph convolution layers with 256, 128, 128, and 64 channels respectively. These graph convolution layers use the SAGEConv algorithm [9] to process their inputs. This algorithm learns two separate weight matrices *W*_1_ and *W*_2_, for node *i* and the neighbors of node *i* respectively. The working of this has been outlined using eq. (2). The method to derive the output from the graph processing block (RiboGL-GNN) is outlined in eq. (3), where 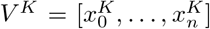 contains either the previous layer node features or the input features when *K* = 0.

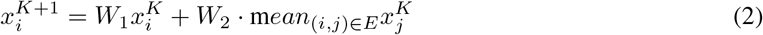

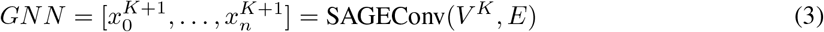

To obtain the final output, the outputs from each layer of the GNN block are concatenated using the Jumping Knowledge module [27]. This helps retain the information from all the layers of the GNN block.
2. **Sequence Processing Block (BiLSTM):** This consists of a 4-layer bi-directional long-short term memory (BiLSTM) model with 128 nodes each. This block was added so that we could potentially augment information by processing the mRNA in terms of a sequence as well.

The overview of the RiboGL model can be found in **Figure 1 (D)**. The model uses learned embeddings from the one-hot encoding of the codon as node features. To train RiboGL, a node-level regression task was performed. For each codon, the model predicted a normalized ribosome density value, which would then be concatenated together to obtain a ribosome profile for the entire gene. The training process was regularized using GraphNorm [5] and Dropout [23] layers. Details about the losses and metrics can be found in Appendix A.2, and the model hyperparameters in Appendix A.3.

### 3.5 Captum Graph Explainer

In order to explain the results on various genes, we studied which neighborhood codons in the graph affected the ribosome density value at the A-site codon in the ground truth ribosome profile. For each gene, all the codons in the ground truth ribosome profile were perturbed using a Captum [13] based explainer employing the “Input X Gradients” [20] algorithm. This method provides attribution values for the chosen codon A-sites with respect to all the nodes, and the edges in the secondary structure graph. This would allow us to understand the relationships of the codons with the sequence and structure neighborhoods. An example of the interpretability output from the Captum Graph Explainer can be found in **Figure 1 (E)**. The height of the orange bars represents the magnitude of the contribution from the codon, and the intensity of the red color for the edges represents the magnitude of their contribution.

## 4. Results

The RiboGL model has a performance of 0.657 in terms of the mean Pearson Correlation Coefficient (PCC) on the mouse liver dataset (section 3.1). Whereas, the state-of-the-art RiboMIMO model which was re-trained and then tested on this dataset has a performance of 0.4459. Therefore, the proposed RiboGL model significantly outperforms the state-of-the-art model by ∼ 21%. This comparison has been outlined in **Figure 2**. Additionally, we compare the RiboGL model with its individual components, the GNN and BiLSTM. The performance of the RiboGL model was ∼ 31% greater than that of the GNN model, but only slightly better than the BiLSTM model. Our hypothesis is that the GNN alone is unable to exploit the full codon context around the A-site due to the limitations of these models with long-range dependencies [1]. The clear difference between the two RNNs, RiboMIMO and BiLSTM, is due to the fact that RiboMIMO does not have a learnable embedding like RiboGL and its components. Lastly, the small increase from the BiLSTM to RiboGL could be due to the fact that structural neighbors only help the model for specific genes, and such a difference is not immediately reflected by our dataset-wise metrics.

**Figure 2.**
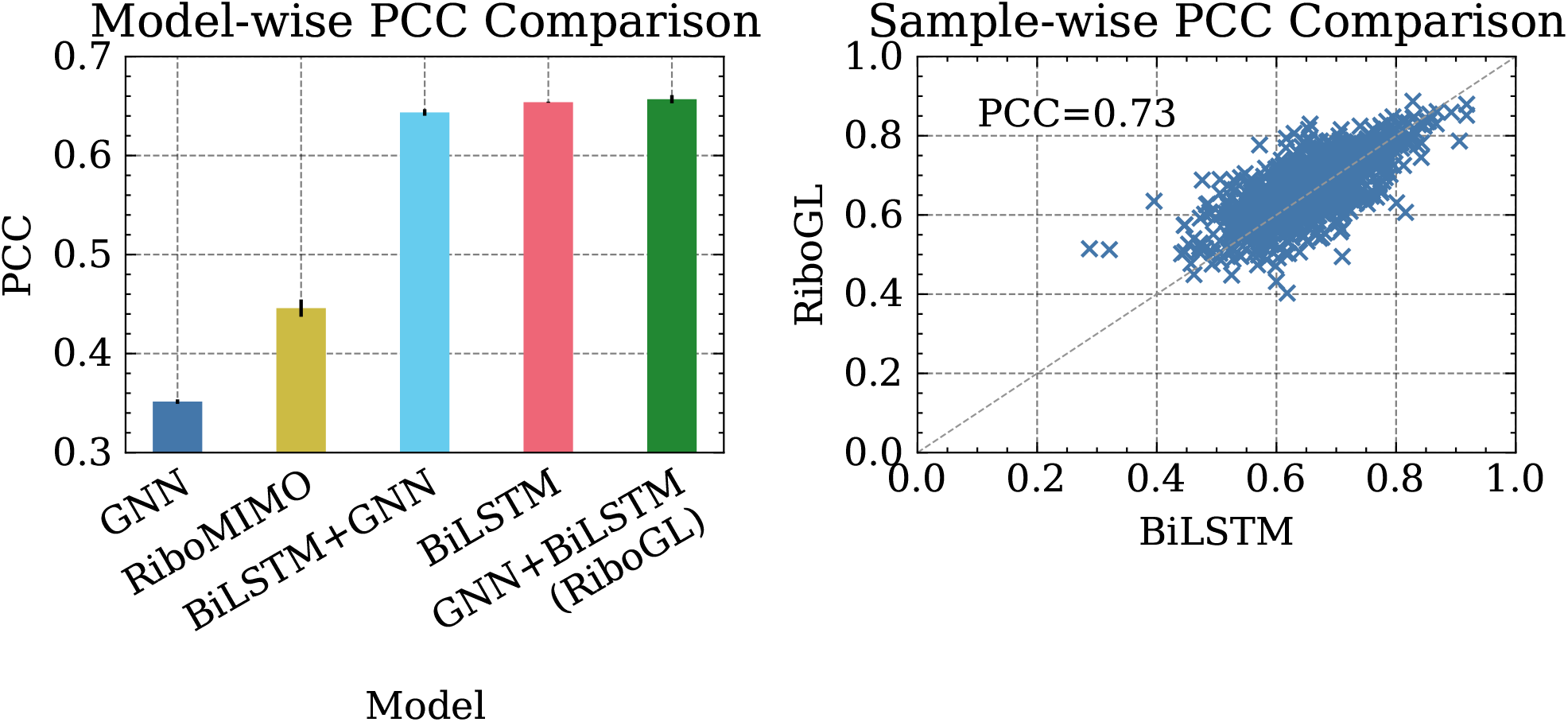
RiboGL comparison with the state-of-the-art RiboMIMO, and the BiLSTM, GNN, and BiLSTM+GNN models. (Left) The individual performances of the models in terms of the Pearson correlation coefficient. Each of them was trained 5 times with different seeds to obtain the mean and standard deviation of the performances. The values can be found in **Table A.1**. (Right) Scatter plot displaying the gene-wise performances of the RiboGL and BiLSTM models on the testing set of mouse liver dataset.

### 4.1 Ablation study on the order of the RiboGL components

To understand the importance of the different design aspects in creating the RiboGL model, we tried two combinations for the order of the graph and sequence processing blocks. The first order with the graph processing block in the beginning, followed by the sequence processing block is the proposed RiboGL model. The second order is the inverse of the first and is named the BiLSTM+GNN model. The proposed RiboGL model improves slightly over the BiLSTM+GNN model (**Figure 2**, and **Table A.1**). This could potentially indicate that the graph processing block at the beginning of the pipeline helps design codon embeddings better fused with the mRNA secondary structure graph information, which the BiLSTM then benefits from later on. Out of the four models developed in this study, GNN, BiLSTM, BiLSTM+GNN, and RiboGL, we choose RiboGL as the proposed model, as it has the highest performance and has the added advantage of being able to conduct interpretability studies on the structure edges.

### 4.2 RiboGL detects long-range codon interactions

The Captum Graph Explainer (section 3.5) was used to study all the genes in the test set of the mouse liver dataset using predictions made by the RiboGL model. The perturbation of all the codons in the true ribosome profile of these genes was plotted to analyze on a global scale the effect of the different codons in the neighborhood of the ribosome A-site. In order to compare the RiboGL model, similar perturbations were calculated for the BiLSTM model. In **Figures 3 (A)** and **(B)**, histograms of the distance to A-site for the top-5 codons that contribute to the codon A-site were plotted for the BiLSTM and RiboGL respectively. We can clearly notice from these figures that the RiboGL model is able to account for long-range codon interactions much better than the BiLSTM model.

**Figure 3.**
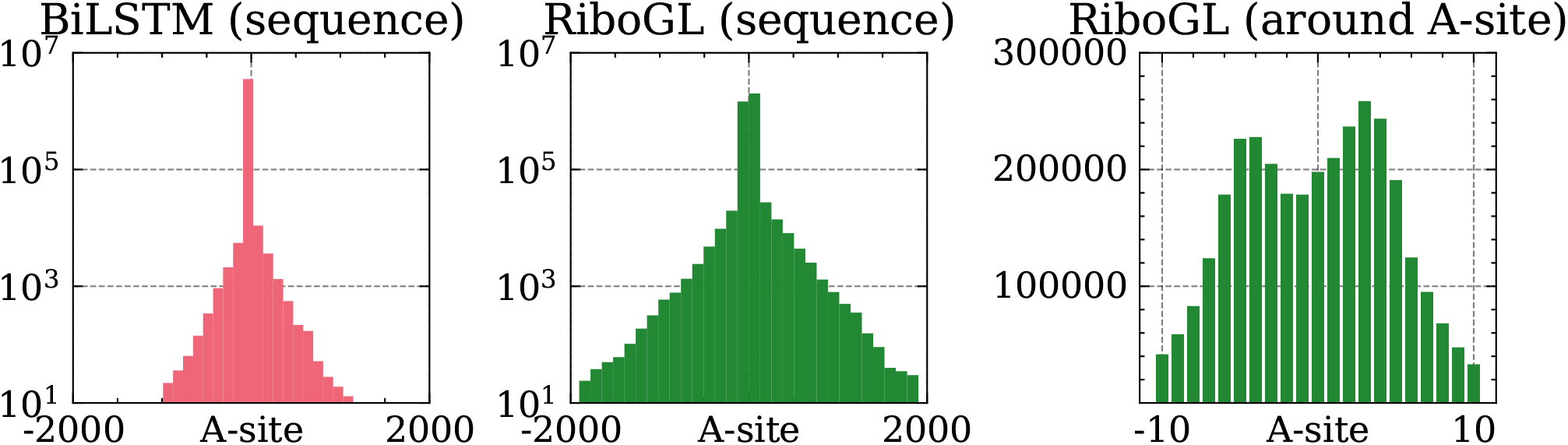
Distribution of distances to A-site of the top-5 codon attributions, for each sequence in the test set. (Left) BiLSTM model. (Center) RiboGL model. (Right) RiboGL model, the distribution was restricted to codons close to the A-site. Comparing BiLSTM against RiboGL, the latter has heavier tails corresponding to long-range interactions.

Additionally, in **Figure 3 (C)** we plot the zoomed-in version of the attributions for the RiboGL model to understand the interactions in the local neighborhood of the A-site. We notice that the codons at the E, P, and A-sites of the ribosome contribute highly to the prediction at the A-site. The codons upstream and downstream from the A-site also contribute to the prediction at the A-site, but their attribution reduces as the distance from the A-site increases. Interestingly, we notice that the codons at positions -4, and +3 have an unusually high attribution to the A-site, which could likely be caused by technical biases that arise from the ribosome profiling process [17, 3].

### 4.3 Structural edge interpretability using RiboGL

The primary advantage of using the RiboGL comes with the fact that we can conduct interpretability studies on the structure edges and learn more about their function. This could not be obtained using the purely sequence-based BiLSTM model as its architecture does not allow it to utilize this structure information. In order to study how the structural edges affect the ribosome count prediction at the A-site we analyze the attributions at an individual gene level. The contributions of the nodes and edges in individual genes in the mouse liver testing set were obtained using the Captum Graph Explainer applied to the proposed RiboGL model. One of the best-performing genes, WD Repeat Domain 43 (“Wdr43”), was chosen to explain this analysis. This gene in humans codes for a protein that acts as a ribosome biogenesis factor [19]. In **Figure 4 (A)**, the predicted and true ribosome profiles of the “Wdr43” gene (in blue and green respectively), along with the global codon-wise contributions (in orange) to the A-site codon at position 623 have been displayed. We notice that the codons in the local neighborhood of position 623 contribute highly to the read counts, but there are also several distant codons, such as position 572, that contribute highly.

**Figure 4.**
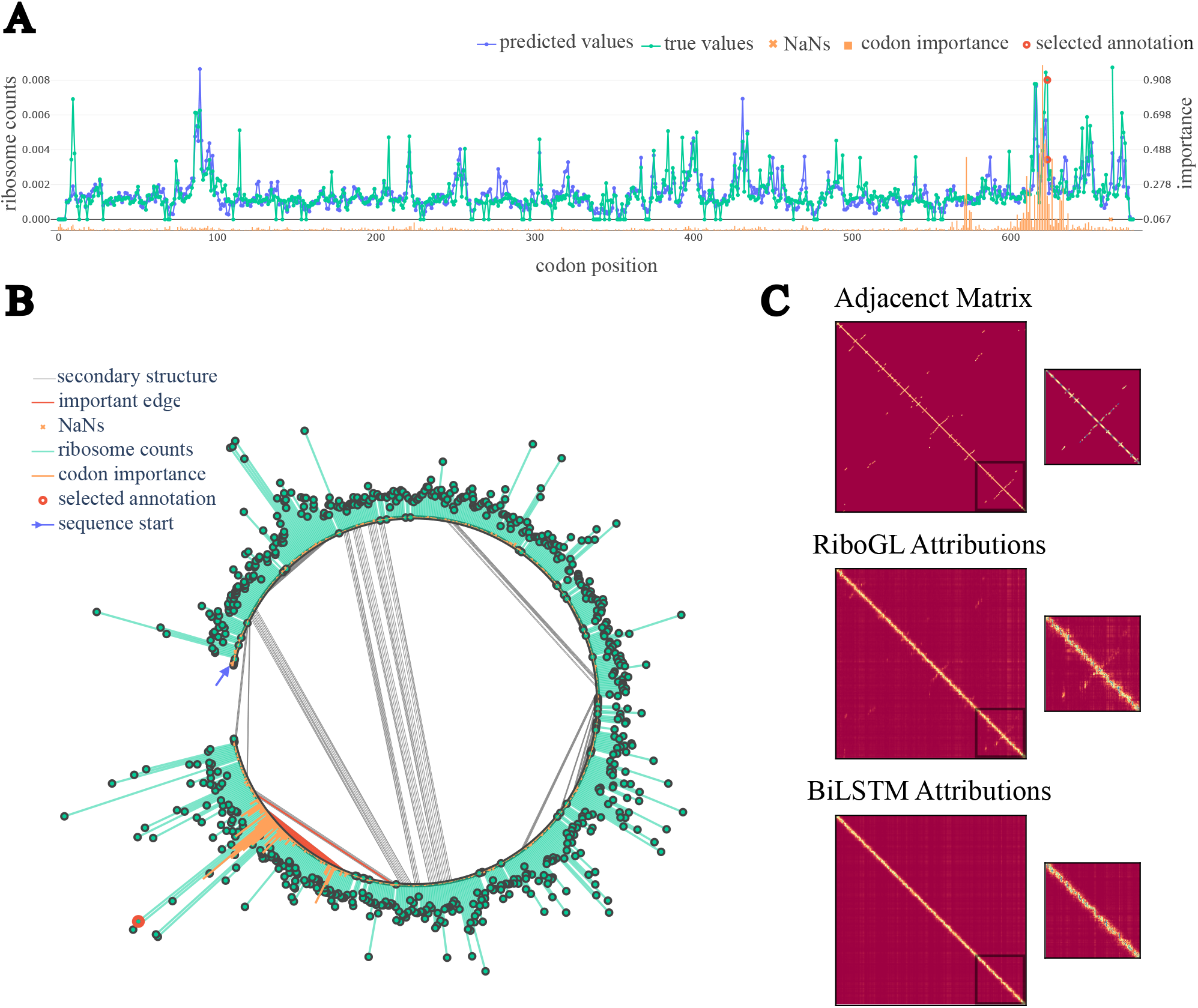
Interpretability analysis using the Captum Graph Explainer built on the RiboGL model on the “Wdr43” gene. (A) True and predicted ribosome profiles on the “Wdr43” gene using the RiboGL model. The codon at position 623 was perturbed using the Captum Graph Explainer, and the node attributions for this position have been displayed. True ribosome profile on the “Wdr43” gene using the RiboGL model. The plot displays the perturbation with all the node and edge attributes of the codon at position 623 which were obtained using the Captum Graph Explainer. The selected codon is the 623 codon (“GAC”), and the long-distance attribution is observed from the codon at position 572 (“CTC”). (C) Three plots for the gene “Wdr43” representing the adjacency matrix of the secondary structure graph, RiboGL attribution matrix, and the BiLSTM attribution matrix. The zoomed-in version of the adjacency matrix plot shows a structure edge at the end of the sequence, which is also seen in the zoomed-in version of the RiboGL attribution plot. However, this is not visible in the zoomed-in version of the BiLSTM attribution plot.

In **Figure 4 (B)**, we check the attributions for the same example using a graph perspective to investigate it further. We notice that the distant codon at position 572 which is connected to the A-site codon at position 623, has a high contribution, this could potentially be because they are connected by structure edges. This contribution of the structure edges has been highlighted in the figure. This is an important advantage of the RiboGL model, as it allows us to understand and model long-range codon interactions and their contributions through the inclusion of structure edges. We further analyze the differences between what the RiboGL and BiLSTM models focus on in **Figure 4 (C)**. We plot the adjacency matrix of the secondary structure graph and study it in the context of the two models’ attributions. The zoomed-in versions of the plots show that the RiboGL attributions pick up on the structure edge links and are able to exploit this information. Whereas the BiLSTM attributions do not show that it is able to pick up on the structure edge information, and hence would not be able to exploit it.

Additionally, to make it easier for one to visualize and study the predictions, and attributions of the RiboGL model, we designed a server that can be launched and hosted locally. This server has all the predictions and attributions for all the 1,397 genes in the testing set of the mouse liver dataset. The server interface allows one to pick any of the genes, and then click on the codon that they are interested in to display the attributions for that particular codon both in the context of the sequence and secondary structure graph, similar to **Figures 4 (A)** and **(B)**. To the best of our knowledge, this server is the first of its kind where one can analyze and interpret the predictions of the machine learning models for ribosome density prediction at a codon resolution. We believe that the scientific community can greatly benefit from this kind of server as it makes the interpretability information highly accessible. Further details on how to run and host this server can be found in our GitHub repository. To provide some additional examples, in **Figures A.3** and **A.4** we reported the plots for sequence and structure level predictions of the top-6 predicted genes, while in **Figure A.5** we provided the plots of selected codon attributions.

### 4.4 RiboGL and LSTM Performance Analysis

We further investigate the RiboGL and the BiLSTM models using a scatter plot of their performances on the testing set samples, which is displayed in **Figure 2 (B)** (0.73 PCC). We can notice from this plot that the distribution is mostly on the diagonal, showing that these models have similar performances on all of the sequences. However, the addition of the GNN on top of the BiLSTM allows us to incorporate and study the structure edges.

## 5 Limitations

The proposed RiboGL model greatly outperforms the state-of-the-art RiboMIMO model and lays the foundations for modeling ribo-seq data as a graph. However, while the example outlined in the discussion section 4.3 showcases the importance of the structure edges, we did not get a definitive result regarding the performance. We highlight that the information coming from structural neighbors might affect only specific genes or codons, and therefore be lost when averaging the metrics across the whole dataset. Moreover, the results could be highly dependent on the characteristics of the data, such as the coverage distribution (see A.1). Investigating additional bigger datasets, such as those belonging to other species, could help clarify the role of structural neighbors in density prediction. Lastly, tools such as “Input X Gradient” should be interpreted with care by taking into consideration their reliability [2]. As a future work, we plan to implement also approaches to increase the robustness of the results, for example by smoothing interpretations [21].

## 6 Conclusion

RiboGL is the first graph-based approach applied to predicting full-length ribosome densities. We showcase the importance of using the physical and measurable aspects of mRNA such as the predicted secondary structure, and how we can process this in terms of a graph. We improved the existing state-of-the-art RNN [24] and combined it with a GNN to make it able to process both the mRNA sequence and secondary structure graph features. We also showed how the graph-based interpretability method can be used to study individual genes and to understand how node and edge attributions affect the prediction at the A-site. In the future, we would like to explore the relationship of ribosome density values with other aspects of the translation, such as the structure of the nascent polypeptide, that can be also modeled by a GNN.

## 7 Acknowledgments

The authors thank the Swiss National Science Foundation (SNSF) for funding this study (SNSF: # 205884). Additionally, we thank Ali Hariri, Maria Boulougouri, Anaïs Haget, and David Neill Asanza for their valuable insights and discussions.

## A. Appendix

### A.1 Preprocessing

The following steps were conducted after obtaining the pre-processed reads from the Ribo-DT pipeline:

1. **Gene-wise Normalization:** The ribo-seq experiments for this dataset were conducted multiple times to obtain 84 replicates. The codon-level ribosome read counts for these individual replicate experiments were averaged within each gene. Only the longest transcript for every gene was chosen, and the others were removed.
2. **Merging:** The gene-wise normalized ribo-seq replicates were merged to obtain one annotation of ribosome densities per gene.
3. **Annotation:** The codons in all the gene sequences were assigned their respective gene-wise read count values and normalized by the average count. The ranges of these counts were reduced by applying the log1p function.
4. **Pruning:** Multiple threshold conditions were applied while choosing to keep a gene in the final dataset. Those conditions were:
  - Long Sequence of NaNs: All of the gene sequences were traversed and if there were contiguous stretches of zero counts longer than 30 codons, these were converted into NaNs. Once the zeros were converted to NaNs, if those genes had more than 5% of their codons annotated with NaNs, they were removed from the dataset.
  - Long Sequence of Zeros: Genes that had a contiguous sequence of zero count values greater than 20 codons in length were removed from the dataset.
  - Percentage of Sequence Annotated (Coverage): Genes that had less than 30% coverage were removed from the dataset. The coverage was defined by the number of codons annotated with a non-zero and non-NaN count value, divided by the number of codons annotated with a non-zero count value.
5. **Dataset Split:** The resulting dataset consisted of 6,988 genes and this was split into training and testing sets. The genes in the dataset were first ordered in descending order of their coverage. Alternating sequences from the top of this list were added into training, and testing sets until the testing set consisted of 20% of the original dataset. The training set consisted of 5,591 genes and the testing set consisted of 1,397 genes. This kind of train-test split was chosen to maintain the quality of the testing set while reducing the redundancy.

#### A.2 Loss and Metrics

To conduct this training process, a multi-term loss function combining the Pearson Correlation Coefficient (PCC) and mean absolute error (MAE) was designed. The equations for the loss function have been outlined below.

The PCC was used to evaluate the performance of the model, and it was calculated between the predicted sequence of ribosome densities and the true sequence of ribosome densities.

1. Mean Absolute Error (MAE or L1):

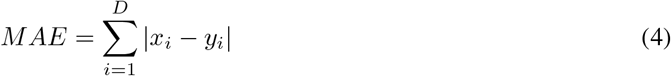
2. Pearson Correlation Coefficient (PCC):

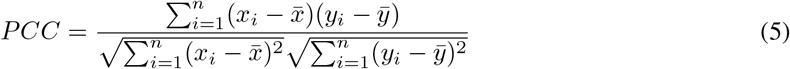
3. RiboGL Loss

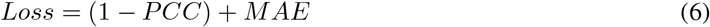

#### A.3 Model Hyperparameters

The RiboGL model was trained for 200 epochs with an early stopping patience of 20 epochs. The AdamW [16] optimizer was used with a learning rate of 1*e* − 2, which was reduced by a factor of 0.1 every 10 epochs of the loss not decreasing. A batch size of 1 and a dropout of 0.3 was used for the training process.

**Figure A.1:**
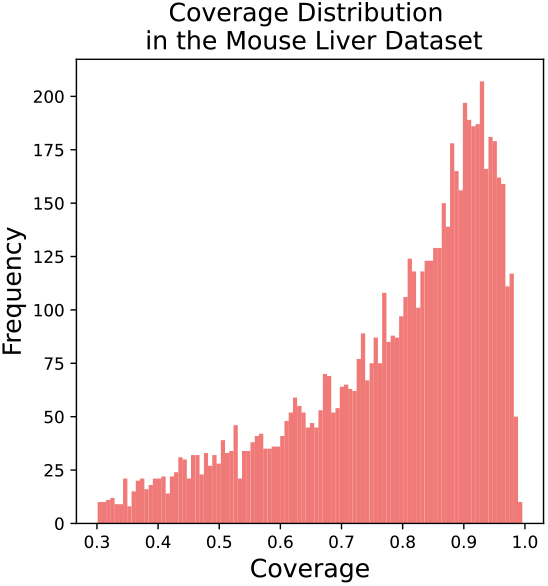
Coverage distribution of the genes in the mouse liver dataset.

**Figure A.2:**
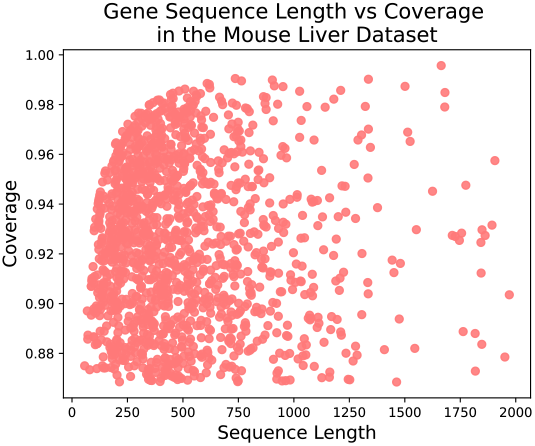
Scatter plot of the coverage and the sequence length of the genes in the mouse liver dataset.

**Table A.1:** Performance comparison. Performance comparison of the 4 configurations of GNN and BiLSTM, against the current state-of-the-art RiboMIMO. Note that to compute MAE for RiboMIMO we normalized the predictions to obtain a probability distribution.

| Model | PCC | MAE |
| --- | --- | --- |
| GNN | $0.351 \pm 0.002$ | $0.032 \pm 0.00006$ |
| RiboMIMO | $0.446 \pm 0.009$ | $0.649 \pm 0.01380$ |
| BiLSTM+GNN | $0.643 \pm 0.003$ | **0.029** $\pm 0.00008$ |
| BiLSTM | $0.654 \pm 0.0006$ | $0.035 \pm 0.00001$ |
| GNN+BiLSTM | **0.657** $\pm 0.0041$ | **0.029** $\pm 0.00013$ |

**Table A.2:** Model size. Number of parameters in each of the models

| Model | No. of Parameters |
| --- | --- |
| GNN | 838,529 |
| GNN + BiLSTM (RiboGL) | 2,746,945 |
| BiLSTM | 1,598,211 |

**Figure A.3:**
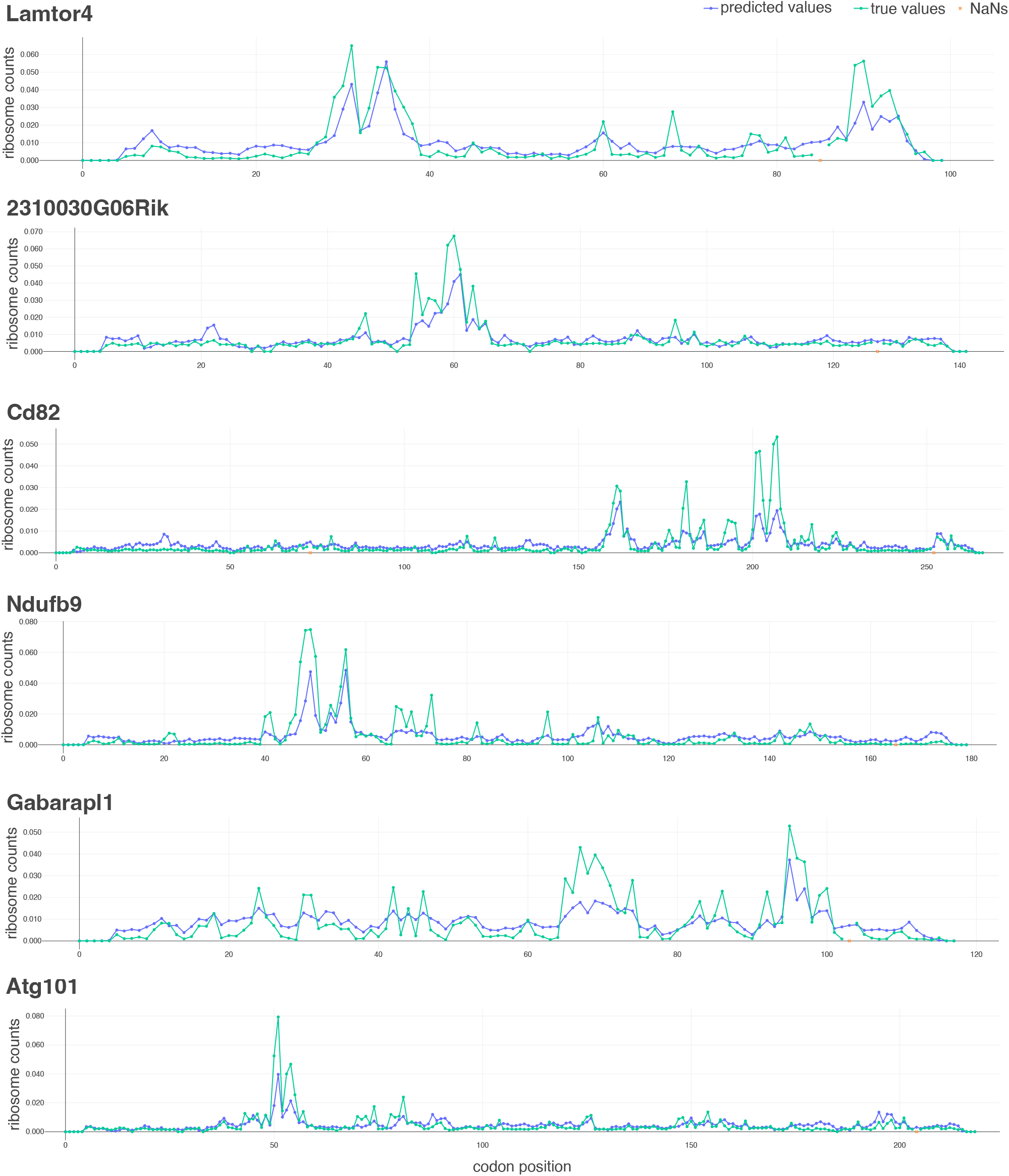
Sequence plot of the top-6 predicted genes by PCC. From top to bottom, the top-6 predicted genes are: Lamtor4 (PCC=0.89), 2310030G06Rik (PCC=0.88), Cd82 (PCC=0.86), Ndufb9 (PCC=0.86), Gabarapl1 (PCC=0.85), and Atg101 (PCC=0.85).

**Figure A.4:**
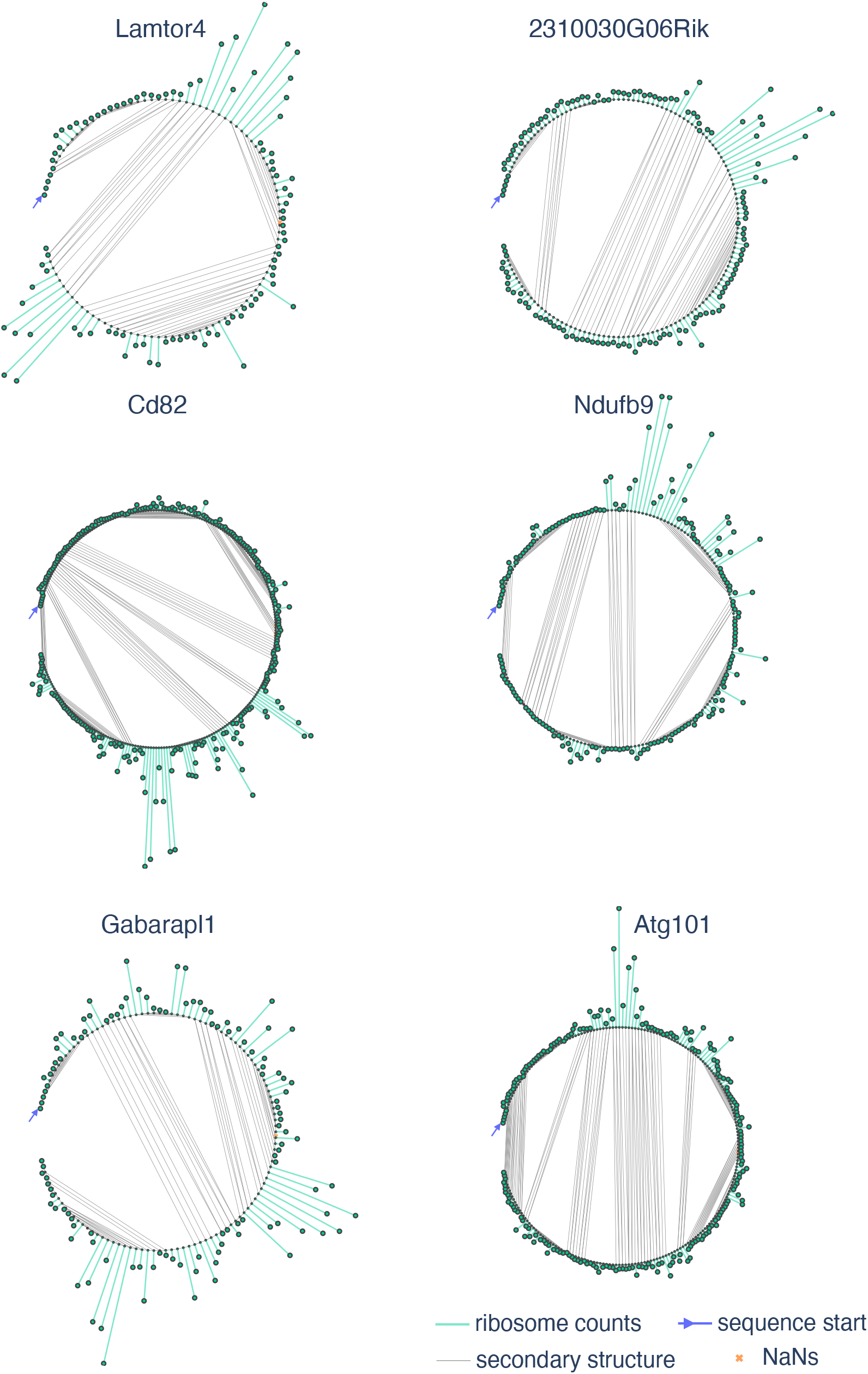
Graph plot of the top-6 predicted genes by PCC. Plots with structural edges of genes included in Appendix Figure A3.

**Figure A.5:**
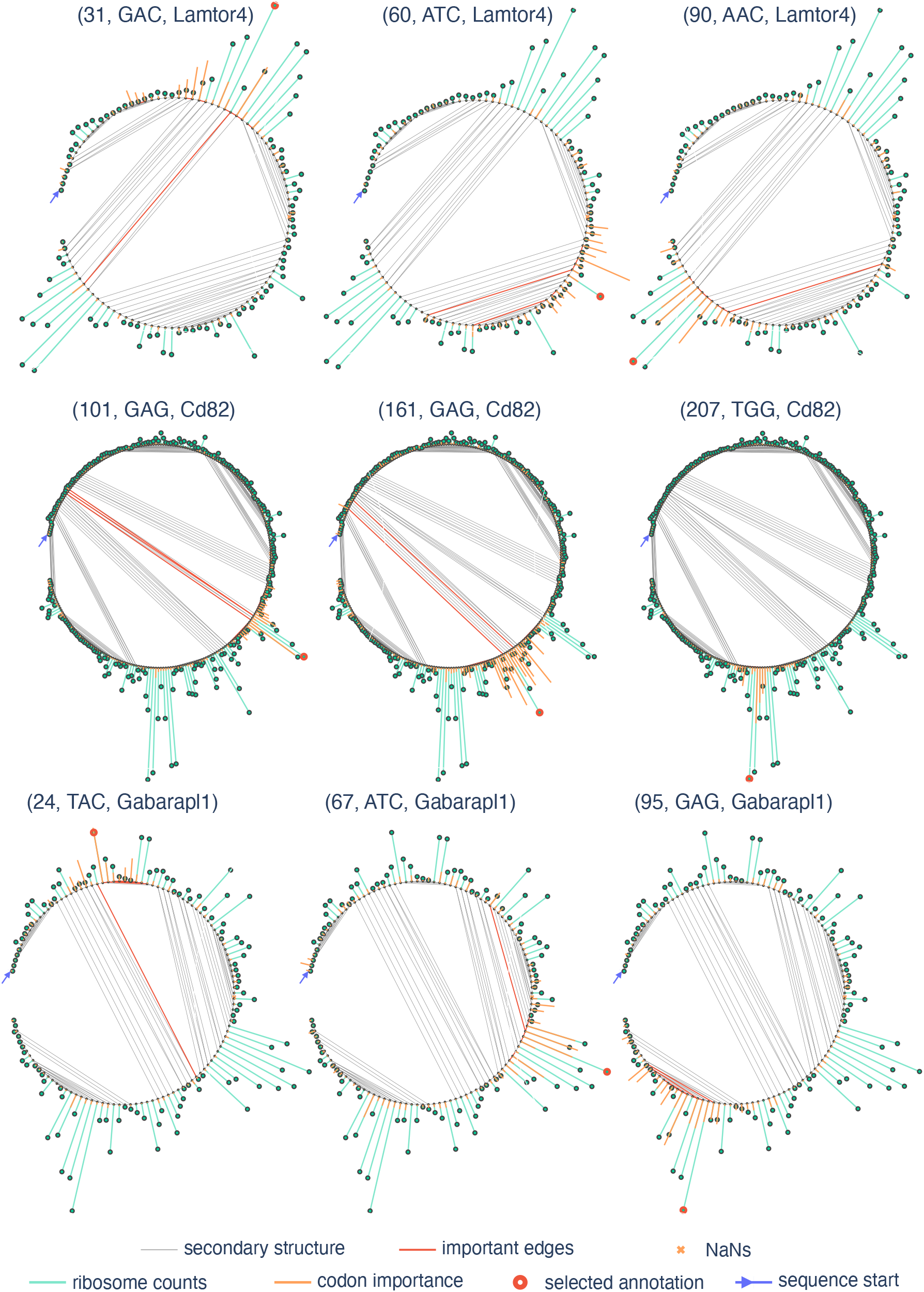
Examples of codon attributions. Visualization of codon attributions for selected genes and codon positions. Each title reports the codon position, the codon name and the gene, respectively.

#### Footnotes

2 https://github.com/vam-sin/ribogl/

3 https://github.com/cgob/codonDT_snakemake

